# An intravenous DNA-binding priming agent protects cell-free DNA and improves the sensitivity of liquid biopsies

**DOI:** 10.1101/2023.01.13.523947

**Authors:** Shervin Tabrizi, Carmen Martin-Alonso, Kan Xiong, Timothy Blewett, Sainetra Sridhar, Zhenyi An, Sahil Patel, Sergio Rodriguez-Aponte, Christopher A. Naranjo, Shih-Ting Wang, Douglas Shea, Todd R. Golub, Sangeeta N. Bhatia, Viktor Adalsteinsson, J. Christopher Love

## Abstract

Blood-based, or “liquid,” biopsies enable minimally invasive diagnostics but have limits on sensitivity due to scarce cell-free DNA (cfDNA). Improvements to sensitivity have primarily relied on enhancing sequencing technology *ex vivo*. Here, we sought to augment the level of circulating tumor DNA (ctDNA) detected in a blood draw by attenuating the clearance of cfDNA *in vivo*. We report a first-in-class intravenous DNA-binding priming agent given 2 hours prior to a blood draw to recover more cfDNA. The DNA-binding antibody minimizes nuclease digestion and organ uptake of cfDNA, decreasing its clearance at 1 hour by over 150-fold. To improve plasma persistence and limit potential immune interactions, we abrogated its Fc-effector function. We found that it protects GC-rich sequences and DNase-hypersensitive sites, which are ordinarily underrepresented in cfDNA. In tumor-bearing mice, priming improved tumor DNA recovery by 19-fold and sensitivity for detecting cancer from 6% to 84%. These results suggest a novel method to enhance the sensitivity of existing DNA-based cancer testing using blood biopsies.

## Main

Liquid biopsies enable non-invasive cancer detection (*1*). Beyond oncology, liquid biopsies also have utility in non-invasive prenatal testing (*2*), infectious disease diagnostics (*3*), and transplant medicine (*4*), with a market value projected to exceed $10 billion in the next decade (*5*). Cell-free DNA (cfDNA) is a major analyte recovered from liquid biopsies. Advances in sequencing methods and related analytical approaches over the past decade have improved performance of tests using liquid biopsies. Yet, despite these advances, sensitivity remains a fundamental challenge. In oncology, ctDNA screening tests only detect 20-40% of stage I tumors and tests for minimal residual disease (MRD) only have 25-50% sensitivity after surgery, when decisions on adjuvant therapy are needed (*6–10*). Detection of resistance mutations for targeted therapy selection can fail in up to 40% of patients with advanced cancer overall, and in up to 75% of patients early on in development of resistance, when resistance mutations are rare in plasma (allele fraction < 1%) (*11–13*).

The quantity of cfDNA in a blood draw is minute. For instance, for a patient with a 1cm tumor (~10^9^ cancer cells), 2-3 blood draws yields only 60 ng of cfDNA. Of this quantity, only ~6 pg (or, one cancer cell equivalent) originates from the tumor (ctDNA) (*14*). This intrinsic scarcity of the primary analyte limits the sensitivity of liquid biopsies, regardless of the technical performance of the *ex vivo* technologies used, creating a significant barrier for the utility of these tests (*15–17*). Collecting larger quantities of blood can only linearly increase the amount of ctDNA recovered and is often not practical (*18, 19*). Some have relied on sampling biofluids other than plasma (*20–23*), but these are often tumor-type specific (and not applicable to cancers without accessible biofluids), do not provide information about distant sites and potential micro-metastatic disease (which is key for relapse monitoring), and often require specialized, expensive, and invasive procedures. Therefore, alternatives to improve the recovery of ctDNA within the standard practices of blood sampling could substantially improve the benefits of ctDNA-based tests.

Cell-free DNA is rapidly cleared from circulation via degradation by circulating nucleases and organ clearance (*10, 14*). We reasoned therefore that delivering a priming agent that could bind and protect cfDNA from clearance a few hours prior to a blood draw could allow recovery of more ctDNA and increase sensitivity for detecting cancer-specific mutations (**Fig. 1A**). We selected monoclonal antibodies (mAbs) as the class of molecules to develop given their persistence in circulation, ease of engineering, established manufacturing processes, and prevalence as biopharmaceuticals (*24–26*). Here we demonstrate a DNA-binding antibody priming agent that increases ctDNA detection by >10-fold in preclinical models. We demonstrate that this agent binds free and histone-bound dsDNA (the components of cfDNA), decreases clearance of dsDNA from plasma, and improves the detection of ctDNA in plasma by up to 19-fold. Together, these results represent the first demonstration of a DNA-binding priming agent for liquid biopsies, with broad implications for the field.

**Fig. 1:**
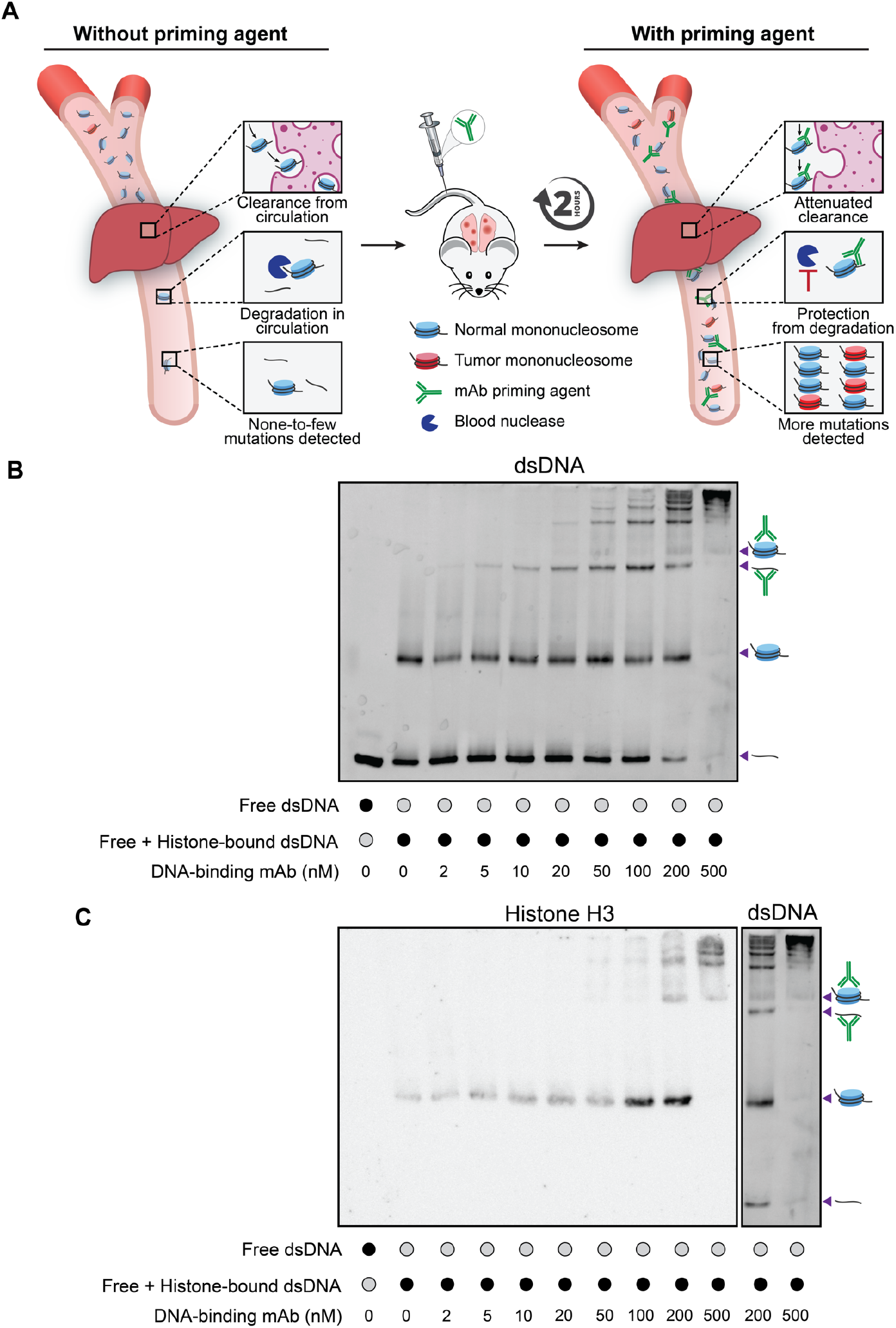
Candidate priming agent binds both free and histone-bound DNA. **(A)** Schematic of approach to protect cfDNA from major clearance pathways of nuclease-mediated degradation and organ clearance via coupling to a DNA-binding priming antibody. **(B)** Electrophoretic Mobility Shift Assay (EMSA) of free dsDNA and histone-bound dsDNA (4ng/μL total dsDNA per lane) with varying concentrations of DNA-binding mAb 35I9 in PBS. **(C)** Immunoblot of the gel in (**B**) for human histone H3. The two right-most lanes from (**B**) are included to facilitate comparison of bands. Relevant bands representing various dsDNA complexes are indicated.

We first tested eight known mouse anti-DNA IgG antibodies for double-stranded DNA (dsDNA) binding activity (**Fig. S1A**). Four demonstrated detectable binding at sub-nanomolar concentrations. Of these four, we selected a mouse IgG2a mAb derived from a NZWxNZB F_1_ lupus prone mouse (35I9) for further investigation. It binds both dsDNA (K_d_ = 90 nM) and ssDNA (K_d_ = 710 nM) (*27*). We explored the interaction of this antibody with free dsDNA in an electrophoretic mobility shift assay (EMSA) and observed shifted bands corresponding to dsDNA-mAb complexes at mAb concentrations above 10nM, with formation of complexes with higher ratios of mAb to dsDNA at higher mAb concentrations (**Fig. S1B**).

Given that the most abundantly observed fragment length of cfDNA is 166bp, corresponding to the size of dsDNA wrapped around a histone octamer with flanking linker dsDNA, cell-free DNA is hypothesized to circulate mostly as mononucleosomes with some free dsDNA, (*28, 29*). To evaluate how our mAb interacted with cfDNA, we next examined its association with a mixture of free and nucleosome-bound dsDNA on EMSA. We observed the appearance of shifted bands with increasing mAb concentration and near disappearance of the free dsDNA and histone-bound dsDNA bands at the highest mAb concentration (500nM), demonstrating the ability of the mAb to bind both free and histone-bound dsDNA (**Fig. 1B**). Consistent with these EMSA results, we observed reduced contrast of isolated mononucleosomes in presence of higher ratios of 35I9 on transmission electron microscopy (TEM) (**Fig. S2).** We also observed the appearance of additional bands on EMSA between those corresponding to discrete ratios of antibody to free dsDNA (**Fig. 1B and Fig. S1B**), and hypothesized that these may represent supershift bands, resulting from binding of mAb to dsDNA that remains histone bound. Immunoblotting of the same gel for human histone H3 confirmed the presence of histones in these supershift bands and revealed multiple H3-positive bands, consistent with binding of more than one mAb to histone-bound dsDNA (**Fig. 1C**). Together these data demonstrate the ability of one or more mAb molecules to interact with both free and histone-bound DNA, the constituent components of cfDNA.

Based on these results *in vitro*, we next asked how this interaction affects the clearance kinetics of cfDNA *in vivo*. We injected a preparation of histone-bound and free dsDNA into Balb/c mice with or without the 35I9 mAb (or an unrelated IgG2a control) and measured levels of the injected dsDNA in plasma via serial blood draws (**Fig. 2A**). The 147-bp Widom601 sequence (W601) was used in the injected preparation and its level in plasma was quantified via qPCR (*30*). After 60 minutes, the absolute quantity of W601 measured was similar between the groups (**Fig. 2B**). This outcome was due to an initial rapid clearance in the mAb-treated groups compared to controls: for example, after 1 minute the quantity of W601 was 1.03 pg/μL with dsDNA alone versus only 0.10 pg/μL with 40μg mAb (P = 0.036). Between 1 minute and 60 minutes, however, clearance of W601 was significantly delayed with mAb treatment. When normalized to 1-minute post-injection levels, 1.9% of W601 remained at 60 minutes with 40μg mAb versus only 0.04% with dsDNA alone (P = 0.008; **Fig. 2C**).

**Fig. 2:**
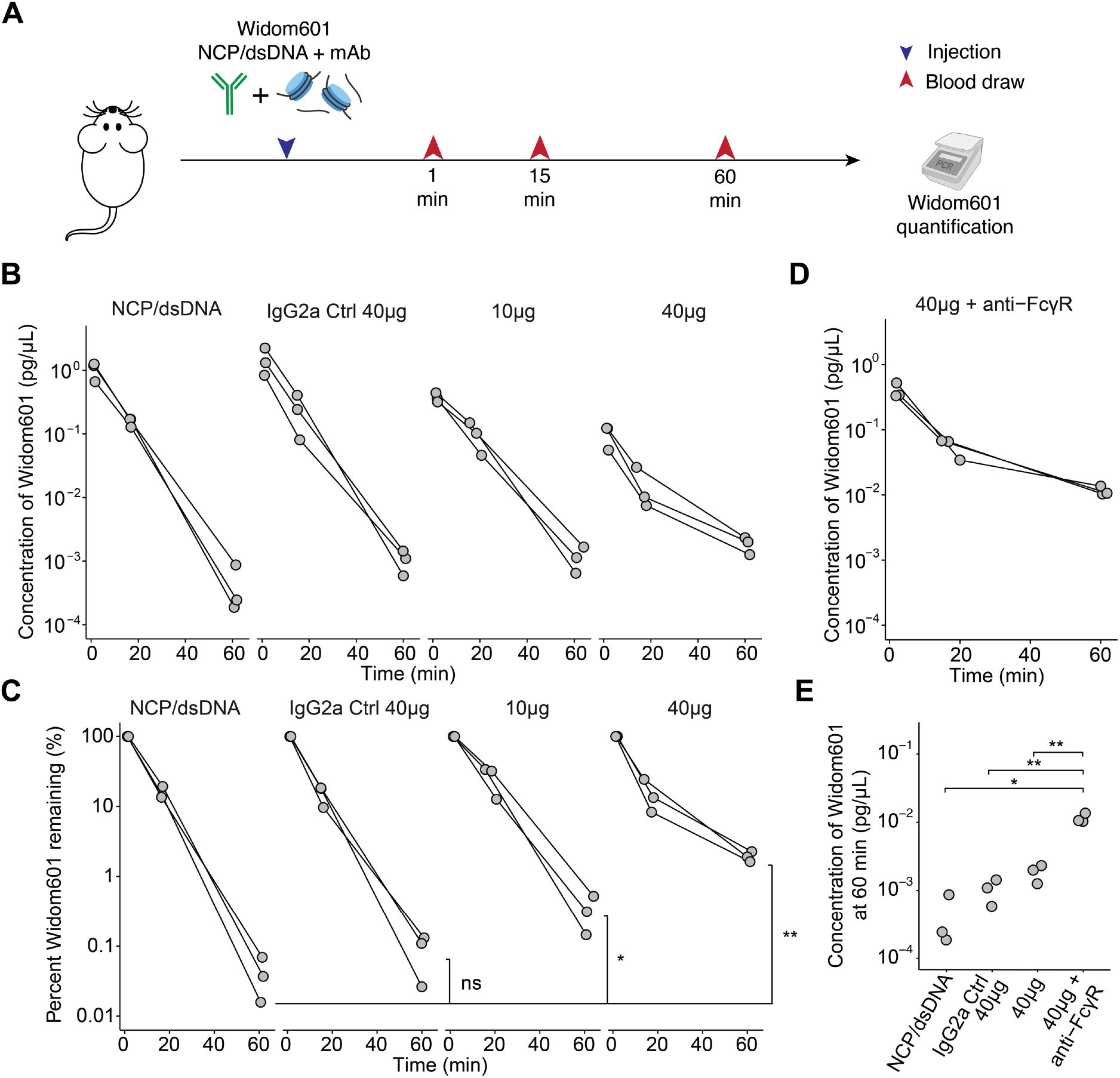
Candidate priming mAb alters the clearance kinetics of circulating dsDNA. **(A)** Experimental approach to evaluate the effect of priming mAb on dsDNA clearance. Nucleosome-bound and free dsDNA with or without mAb were co-injected followed by serial blood draws and qPCR quantification of the W601 sequence in plasma. **(B)** Absolute quantification of plasma W601 levels over time after injection of W601 preparation alone, co-injection with an unrelated IgG2a antibody (IgG2a Ctrl), or with various amounts of DNA-binding antibody 35I9 (10μg or 40μg). **(C)** Percent clearance of W601 over time, normalized by 1-minute levels. (**D)** Effect of anti-FcγRI (20μg) and anti-FcγRII/III (40μg) co-injection on clearance kinetics on W601 when injected with 40μg of DNA-binding antibody 35I9. **(E)** Concentration of W601 detected in plasma 60 minutes after injection. **B-E**, data for n=3 mice per condition. ns = not significant, * P < 0.05, ** P < 0.01; one-way ANOVA.

Given the near immediate clearance of a significant fraction of W601 dsDNA when administered with 35I9, we hypothesized that some of the W601 dsDNA complexed with mAb was rapidly sequestered and cleared from plasma via the mononuclear-phagocyte system (MPS), which can bind and phagocytose IgG-antigen complexes (*31*). This effect could relate to some of the larger complexes observed *in vitro* (**Fig. 1B**) and would be consistent with prior observations of biphasic clearance of sheared gDNA complexed with IgG (*32*). Fc-γ-receptors (FcγR) on macrophages are central to MPS-mediated clearance of complexes (*33–35*). To evaluate the role of FcγR in W601 clearance in the presence of the mAb, we co-injected the W601-mAb preparation with antibodies blocking mouse FcγRI and FcγRII/III. FcγR blockade yielded higher W601 at one-minute post-injection compared to 40 μg mAb alone (0.4 pg/μL versus 0.1 pg/μL, P = 0.029), partially reversing the observed early clearance of W601. At 60 minutes, FcγR blockade also resulted in higher W601 levels compared to 40μg mAb alone, NCP/dsDNA alone, and IgG2a isotype control (P=0.007 for all comparisons, **Fig. 2D-E**). Together, these results suggested that administration of DNA-specific mAbs can delay the clearance of dsDNA from plasma, but that FcγR-mediated clearance of dsDNA bound to mAb reduces the benefits for prolonged stabilization of the dsDNA.

Having identified FcγR as an important mediator of the early clearance of circulating DNA bound to our priming agent, we next sought to overcome this limitation by engineering the Fc-domain of our priming agent to abrogate FcγR interactions. We selected three sets of mutations known to disrupt FcγR binding (in addition to other effector functions such as complement fixation) (*36*) – aglycosylated N297A (denoted aST2) (*37–39*), L234A/L235A/P329G (denoted aST3) (*40*), and D265A (denoted aST5) (*41, 42*) (**Fig. 3A and Fig. S3**). We confirmed that the engineered mAbs still bound dsDNA (**Fig. S4**). We then tested the effect of our Fc-engineered mAbs on circulating DNA clearance by injecting histone-bound and free W601 dsDNA with our mAbs or an IgG2a control. We included aST1, which did not contain any Fc mutations but was prepared using the same process as our engineered variants, as an additional control (the mAb used in the experiments in **Fig. 2** was also included, with or without co-injection with FcγR-blocking antibodies). Among our engineered variants, aST3 (carrying the L234A/L235A/P329G mutations) demonstrated the highest W601 recovery at 60 minutes (0.64 pg/μL versus 0.0041 pg/μL with IgG2a isotype and 0.099 pg/μL with WT + anti-FcγR; P = 0.009 and P=0.007 respectively; **Fig. 3B**).

**Fig. 3:**
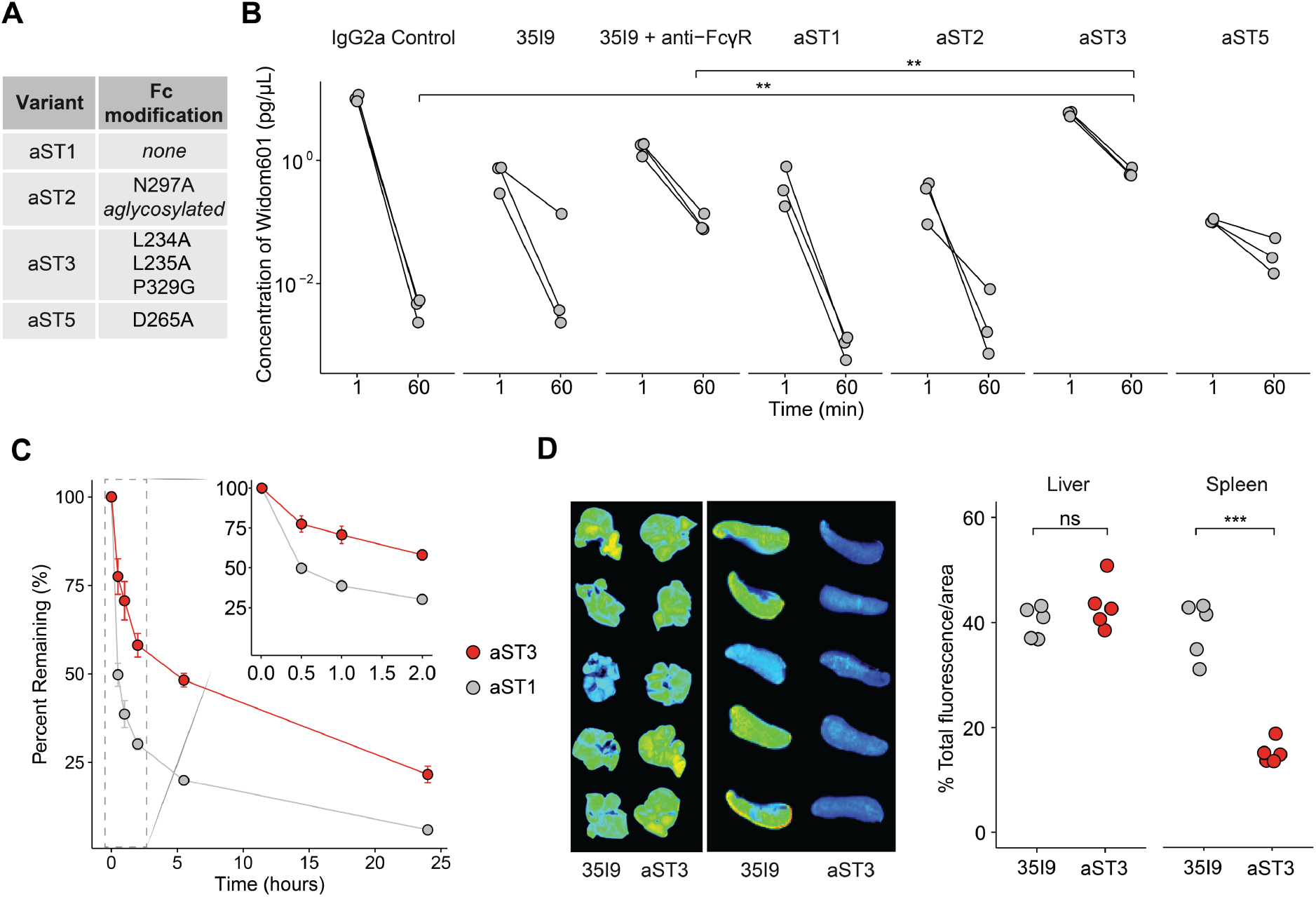
Fc-engineered priming mAb aST3 decreases clearance of circulating dsDNA. **(A)** Engineered variants with Fc-mutations to disrupt interactions with FcγR. **(B)** Concentration of W601 in plasma at 1-minute and 60minutes after injection with various antibodies. IgG2a Control is an unrelated mouse IgG2a mAb. n=3 mice per treatment group. ** P<0.01, one-way ANOVA. **(C)** Pharmacokinetics of aST1 and aST3 labeled with AQuora750 in plasma. Each point represents the mean +/- s.e.m. of n=5 mice per group. **(D)** Biodistribution of aST3 or WT antibody in liver and spleen 1-hour after administration. Individual data shown for n = 5 mice per group. ns = not significant, *** P < 0.001, one-way ANOVA.

To further characterize the effect of this modification on the pharmacokinetics (PK) of our priming agent, we labeled aST3 and the native variant aST1 with a fluorophore and tracked their kinetics in plasma. A difference in clearance was apparent at 30 minutes post-injection and continued to persist, with higher levels of aST3 in plasma observed at all time-points (e.g. 78% of aST3 remaining versus 50% of aST1 at 30 minutes, P = 0.003; **Fig. 3C**). Consistent with this result, we observed less uptake of aST3 in the spleen (**Fig. 3D**). The area-corrected amount of uptake was similar in the liver, the other major organ in MPS, which may be due to differences in the clearance of complexes by the two organs (*43*). Together, these data suggest that the Fc-engineered variant aST3 with abrogated FcγR binding persists longer in plasma and decreases the clearance of circulating DNA compared to the native variant and an IgG2a control.

### Priming improves recovery of normally depleted cfDNA regions

We next explored the effect of our priming agent on endogenous cfDNA. To establish an *in vivo* model of cfDNA and ctDNA, we inoculated Balb/c mice with the luciferized Luc-MC26 murine colon adenocarcinoma cell line to establish lung tumors (**Fig. 4A**). To enable full characterization of unique cfDNA molecules, we designed and validated probes against 1,822 sites in the mouse genome (**Table S1**) using methods previously described (*6*), and used deep sequencing with unique molecular identifiers (UMIs) combined with probe hybrid capture to fully catalogue the unique cfDNA molecules overlapping these 1,822 sites. These sites each overlapped an SNV found in Luc-MC26 but not in normal mouse gDNA, which also enable detection of Luc-MC26 ctDNA in plasma. We injected mice with an IgG2a control or different doses of aST3 and collected blood at two hours post injection. With priming, we observed over ten-fold higher duplex depth per site (e.g. median duplex depth of 863 with 8.0 mg/kg of aST3 versus 57.5 for IgG2a control), which was also consistent with higher cfDNA concentration in plasma (**Fig. S5A,B**).

**Fig. 4:**
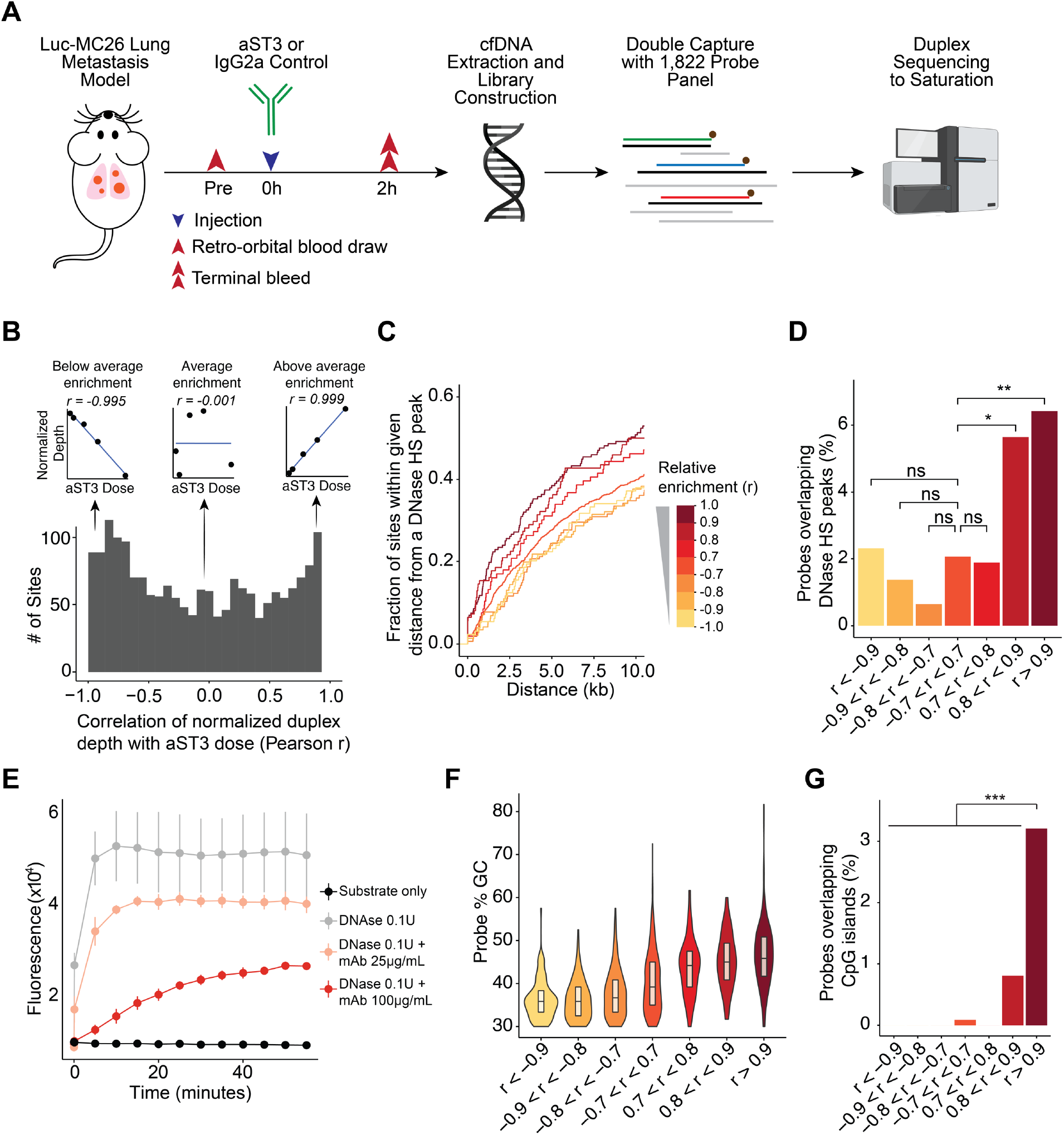
DNase hypersensitivity and GC content are associated with higher relative enrichment from aST3. **(A)** Experimental approach for testing effect of priming on cfDNA in plasma of Luc-MC26 tumor-bearing mice. (**B)** Distribution of Pearson r correlation coefficients between aST3 dose and normalized duplex depth for 1,822 sites. Inserts show scatterplots of normalized depth versus aST3 dose for three sample sites exhibiting less than average, average or more than average enrichment (from left to right). **(C)** Proximity of 1,822 tracked sites to leukocyte/myeloid DNase hypersensitivity peaks in mouse ENCODE, grouped by relative enrichment with higher aST3 doses. **(D)** Fraction of 1,822 probes overlapping leukocyte/myeloid DNase hypersensitivity peaks in mouse ENCODE within each relative enrichment group. **(E)** Fluorescence signal of DNA substrate carrying a hexachlorofluorescein dye on one end and a dark quencher on the other, with or without DNase and mAb 35I9. Digestion of the linking DNA substrate by DNase results in separation of fluorophore and quencher and emission of fluorescent signal. Fluorescence was monitored at 37°C for 1h after addition of 0.1U DNase I. Each point represents the mean +/- s.e.m. of n=3 wells per condition. **(F)** Correlation of GC content of 1,822 probes with the relative enrichment/depletion of the targeted sites with aST3 treatment. Boxplots represent median and interquartile range. **(G)** Fraction of 1,822 probes overlapping CpG islands within each relative enrichment group. All panels refer to experiment with n=6 mice per group. ns = not significant, * P<0.05, ** P<0.01, *** P<0.001; Fisher’s exact test.

Although all sites had higher duplex depth with priming, we asked whether certain sites within our panel benefited more than others from priming. Identifying features of sites that benefit more from priming could provide insights to how our priming agent interacts with cfDNA *in vivo*. To identify sites that were relatively more enriched with use of aST3, we first normalized the duplex depth per site for each dose group. We reasoned that sites that experience the mean overall improvement in duplex depth with aST3 would retain the same normalized depth in all dose groups, whereas those that are enriched relative to the mean would increase in normalized depth with higher doses. We computed the Pearson correlation between aST3 dose and normalized depth to quantify this relationship (**Fig. 4B**) and observed higher normalized depth with higher aST3 doses for positively correlated sites and lower normalized depth for negatively correlated sites (**Fig. S6**). These data suggest that our metric can identify the subsets of sites that are more relatively enriched with our priming agent.

We then explored what features distinguish the relatively enriched sites. Fragmentation of cfDNA by nucleases results in depletion of DNase-sensitive regions of the genome (corresponding to open chromatin) in cfDNA (*28*). Proteins bound to dsDNA can shield DNA from nuclease digestion, which has been utilized to map chromatin accessibility and transcription factor binding sites (*44, 45*). We hypothesized that binding of aST3 to cfDNA could similarly protect cfDNA from nuclease degradation, leading to preferential protection and accumulation of sites that are normally prone to degradation and are depleted in cfDNA. To test this possibility, we collated 13 out of 15 available leukocyte and myeloid DNase-seq hypersensitivity (DNase HS) peak datasets from mouse ENCODE (datasets ENCSR000CMQ and ENCSR000CNP were excluded due to extremely low read-depth), resulting in 1,042,312 peaks (*46*). These datasets were chosen because 85% of cfDNA originates from white blood cells and erythroid progenitors (*47*). We examined the proximity of the sites to DNase HS peaks, and found that sites with the highest relative enrichment after aST3 also tended to be closer to DNase HS peaks (**Fig. 4C**). Consistent with this observation, 6.4% of sites with the highest relative enrichment (r > 0.9) and 5.7% of those with 0.8 < r < 0.9 directly overlapped DNase HS peaks versus 2.1% of those with lower relative enrichment (−0.7 < r < 0.7), P = 0.002 and P = 0.025 respectively, Fisher’s exact test (**Fig. 4D**). To directly test whether our mAb can inhibit nuclease degradation of dsDNA, we incubated our mAb with a free DNA substrate labeled on one end with hexachlorofluorescein and on the other end with a dark quencher, such that digestion of the linking DNA results in a fluorescent signal. In the presence of increasing concentrations of our mAb, we observed diminished signal upon addition of 0.1U DNase I (**Fig. 4E**). These results suggest that our priming agent can protect dsDNA from nuclease digestion, and that *in vivo*, this is associated with higher relative enrichment of DNase hypersensitive sites in cfDNA.

Many antibodies that bind dsDNA preferentially bind to GC-rich regions (*48–52*). We asked whether our cfDNA data reflects this binding preference. We compared the GC-content of our 120-bp probes versus the relative enrichment of the site captured by each probe with varying doses of aST3, and noted higher GC-content at sites with higher relative enrichment (P = 2.1×10^-76^, Wald test; **Fig. 4F**). Given that CpG islands are especially enriched in GC content, we also looked at overlap of our sites with CpG islands in the mouse genome and found that 3.2% of the sites with highest relative enrichment (r > 0.9) overlapped CpG islands versus only 0.1% of all other sites (P = 1.7×10^-5^, Fisher’s exact test; **Fig. 4G**). Although CpG islands are important regulatory elements and therefore overlap DNase HS sites (*53*), we do not believe that the observed associations with DNase HS sites and GC content are explained by the effect on CpG islands alone, given that sites with lower relative enrichment (0.7 < r < 0.9) and without significant overlap with CpG islands also exhibit closer proximity to DNase HS sites (**Fig. 4C**) and higher GC content (**Fig. 4F**). Rather, CpG islands likely represent an especially enriched subset of sites that combine the effects of both higher affinity to GC-rich regions and the relative protection of DNase HS regions mediated by our priming agent. These results imply that subsets of cfDNA fragments may be preferentially enriched based on the binding affinity of the priming agent to particular sequences.

We next asked whether aST3 could improve the detection of ctDNA. The 1822 sites captured by our panel each overlapped a tumor-specific SNV found in the Luc-MC26 cell line, enabling ctDNA detection. Tumor mutations were detected from our hybrid-capture duplex sequencing data as previously described (*6*). We used pre-injection blood volumes to confirm that all groups had similar tumor fractions, and used bioluminescence imaging to confirm that all groups had similar tumor burdens (**Fig. S7A,B**).

We observed consistently higher levels of mutant molecules per μL of plasma with aST3 compared to IgG2a control (**Fig. 5A**), with a dose dependent improvement between 0.5mg/kg and 4.0mg/kg. The maximum effect we observed was at a dose of 4.0mg/kg, with a median of 16.4 mutant molecules/μL compared to 0.86 mutant molecules/μL with the IgG2a isotype (P = 0.009), a 19-fold improvement. Tumor burden did not explain this difference, as tumor burden was generally comparable between the two groups, and if anything, two mice in the aST3 4.0mg/kg group had burdens (total flux 0.78×10^8^ p/s and 2.9×10^8^ p/s) well below the range in the control group (total flux 10.1-44.3×10^8^ p/s), **Fig. 5B**. We also observed detection of a higher fraction of our 1,822 SNV panel with priming (e.g. median 77% of sites detected with 4.0mg/kg versus 15% detected with IgG2a isotype, P = 8.5×10^-6^), **Fig. 5C**. The benefit for the 2.0, 4.0 and 8.0 mg/kg doses was similar: all improved ctDNA recovery by at least 10-fold. No difference in tumor fraction was observed between the groups post-injection (**Fig. S8).**

**Fig. 5:**
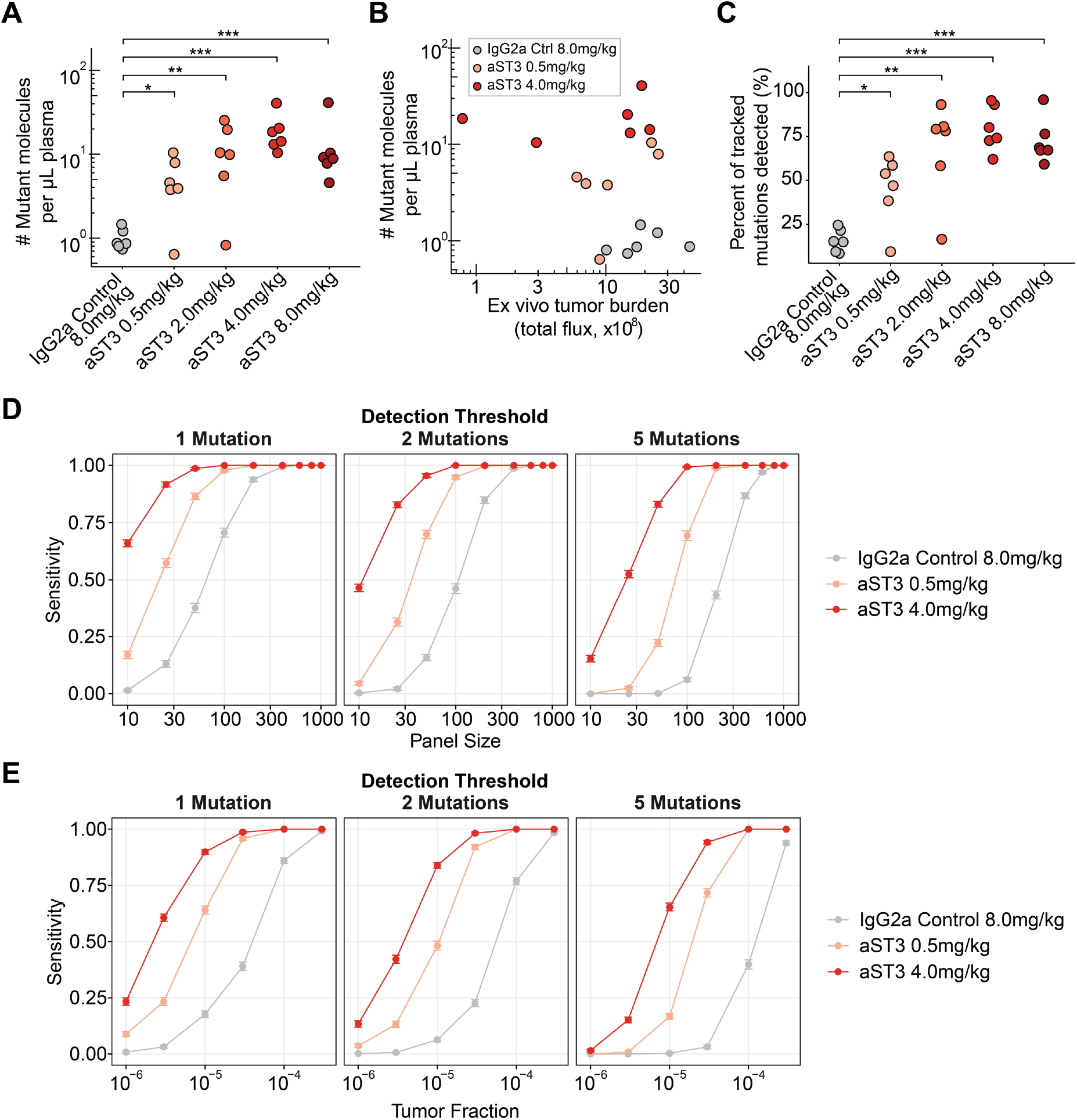
Engineered priming mAb aST3 improves circulating tumor DNA recovery. **(A)** Concentration of tumor-derived, mutant cell-free DNA molecules in plasma after injection of control mAb or various doses of aST3. n=6 mice per group. (**B)** Concentration of tumor-derived, mutant cell-free DNA molecules in plasma after injection of control mAb or various doses of aST3 versus measured ex vivo tumor burden. **(C)** Percentage of sites from a 1,822 SNV panel detected in plasma with control mAb or various doses of aST3. **(D)** Sensitivity for detection of ctDNA in plasma with or without priming agent under various panel sizes and detection thresholds based on down-sampling from full panel. Each point is mean +/- s.e.m. of 100 replicates. **(E) S**ensitivity for detection of ctDNA in mouse plasma with an 1822-probe panel at lower tumor fractions based on binomial down-sampling of mutant molecules. Each point is mean +/- s.e.m. of 100 replicates. IgG2a Control: an unrelated mouse IgG2a antibody. * P<0.05, ** P<0.01, *** P<0.001; one-way ANOVA.

In this experiment, we used a large SNV panel and mice with large tumors to ensure robust ctDNA detection in all groups and enable comparisons between the groups. These are not, however, representative of typical conditions for ctDNA assays in early detection and MRD settings, where ctDNA is used for detection of microscopic disease and monitoring for relapse. Furthermore, current assays typically have much smaller panels, ranging from a single mutation to a few dozen for many commercial assays, and a hundred to a few hundred for some pre-clinical assays (*6, 7, 54, 55*). To determine the effect of our priming agent on the sensitivity of ctDNA assays under conditions for early detection, we first examined the effect with typical panel sizes below the 1,822-SNV panel used in this work. We down-sampled our panel to smaller panel sizes ranging from 10-1000 SNVs and computed sensitivity within each dose group as the mean sensitivity across 100 replicates at each panel size. We have previously used a detection threshold of 2 mutations in clinical samples to call a sample ctDNA-positive (*6*), but included a range of detection thresholds here, recognizing that different clinical assays will have different thresholds and there is no established threshold for preclinical models of ctDNA. Across a wide range of panel sizes and detection thresholds, we consistently observed superior sensitivity with our priming agent (**Fig. 5D**). Using a 2-mutation detection threshold for panel sizes ranging from 10-100 SNVs, sensitivity in the aST3 4.0mg/kg cohort was 47% to 100% versus 3% to 46% in the IgG2a control cohort. This observation suggests that our priming agent can significantly improve the sensitivity of liquid biopsies at panel sizes typical for current commercial tests.

We next evaluated the effect of our priming agent on ctDNA assay sensitivity at lower ctDNA levels. Low fractional content of tumor DNA in plasma (on the order of 1-10 parts per million (ppm)) is typical for early detection settings and is correlated with lower, often microscopic, disease burden (*14, 56*). We first verified that the same correlation between tumor fraction and disease burden held for our samples (**Fig. S9A**). We also observed that the distribution of the number of mutant molecules is recapitulated via a binomial sample using the global tumor fraction for each sample and the total number of molecules at each site (**Fig. S9B-D and Materials and Methods**). To estimate the sensitivity of our ctDNA assay with or without the priming agent at lower tumor fraction, we generated distributions of mutant molecules using binomial sampling from the total duplex depth at each site, but at lower tumor fractions compared to that observed in each sample (**Materials and Methods**). We also tested varying panel sizes as described above. Use of our priming agent resulted in similar sensitivity at approximately 10-fold lower tumor fraction (**Fig. 5E, Fig. S10**). Using a 2-mutation detection threshold, the sensitivity of ctDNA testing improved from 6% to 84% for a 1822-mutation panel at a tumor fraction of 1/100,000 (**Fig. 5E**). The mean plasma volume in our samples was 0.33mL (s.d. 0.09mL), which is approximately one-tenth of the ~4mL of plasma collected in a patient blood draw. With ten-fold higher plasma volume, we would expect a similar improvement in sensitivity at tumor fractions of ~1 ppm. These results suggest that aST3 can significantly improve the sensitivity of ctDNA assays in early detection and monitoring applications.

Despite the commercial use of liquid biopsies for cancer diagnostics today, limited sensitivity remains a major barrier in multiple clinical settings. Here we have developed a DNA-binding priming agent that improves the recovery of ctDNA by 19-fold, significantly boosting the sensitivity of ctDNA assays (**Fig. 5D,E**). This agent binds both free and histone-bound dsDNA and decreases the clearance rate of dsDNA from plasma. We further demonstrate that this agent preferentially enriches cfDNA fragments prone to nuclease degradation and rich in GC content. This work establishes a preclinical proof-of-concept for using a priming agent to boost the sensitivity of liquid biopsy-based diagnostics.

In contrast to other approaches to increase quantities of ctDNA that rely on local sampling procedures (*20–23*), use of a priming agent preserves the use of plasma cfDNA and the advantages it confers, including sampling from all sites that shed DNA into plasma (such as potential micro-metastatic sites) and using a blood draw rather than specialized or invasive procedures. Furthermore, the 19-fold improvement in ctDNA recovery is more than can be reasonably accomplished through routine collection of a larger blood volume (except plasmapheresis, which is too invasive and costly for routine diagnostic use). The use of a priming agent could be incorporated into current collection strategies which involve drawing multiple blood tubes to maximize sensitivity. Although we focused on ctDNA – specifically somatic mutation detection – the use of priming agents could, in principle, be extended to other analytes in liquid biopsies where sensitivity remains a challenge.

Further optimization and development of our technology is needed prior to clinical translation. The pharmacokinetics of cfDNA and the priming agent may differ in larger mammals, which may require changes in dose or formulation. Furthermore, although no acute reactions were observed following priming, investigation of tolerability and toxicity in larger mammals is needed. DNA-binding antibodies have been associated with autoimmune conditions such as systemic lupus erythematosus (SLE), and a subset may have pathogenic roles mediated by complement fixation, cross-reactivity, and immune activation (*57–59*). In healthy mice, however, prior studies suggest that DNA-binding mAb doses well above those used in this study (e.g. cumulative dose of 80 mg/kg over 4 weeks(*60*)) are required to observe renal side-effects, and γ^-/-^ mice lacking FcγR signaling are protected against autoantibody-mediated renal injury (*61*). Based on these reports, we posit that an engineered priming mAb that lacks Fc-effector function could achieve appropriate safety profiles at the doses used in this study. Finally, to be practical for routine clinical use, this agent will need to be administered within a timeframe reasonable for a clinic visit. Although biologics have typically required an infusion over 0.5-1 hour, other methods of administration including an intravenous push in less than one minute(*62*) and subcutaneous depots(*63, 64*) have been established for some mAbs. Optimization of dose and delivery routes for this priming agent will be important for clinical use.

In this work, we pursued a strategy of coupling cfDNA with a protective molecule to inhibit clearance. In a companion paper we demonstrate an orthogonal strategy of inhibiting the cells involved in uptake of cell-free DNA from plasma using a nanoparticle-based approach (manuscript #adf2341). Together, these two approaches demonstrate that multiple, potentially synergistic, priming approaches could be used to improve the recovery of cfDNA. We envision that priming agents may eventually play a similar role to iodinated and gadolinium contrast agents currently in routine clinical use in radiology, but to enhance diagnostic sensitivity of liquid biopsies.

In summary, we have demonstrated here an approach to overcome a key barrier in liquid biopsies. Use of priming agents could significantly boost the sensitivity of liquid biopsies and enable the numerous promising applications of the technology in early cancer detection, detection of MRD, treatment response monitoring, decisions on therapy, and in multiple fields beyond oncology. Our antibody-based platform also opens further engineering opportunities to enrich subsets of cfDNA fragments for specific applications.

## Materials and Methods

### DNA-binding antibodies and ELISA

Mouse IgG antibodies against DNA were obtained from commercial vendors – 404 (cat. sc-66081, Santa Cruz Biotechnology), 1.BB.27 (cat. sc-73064, Santa Cruz Biotechnology), 3H12 (cat. sc-73065, Santa Cruz Biotechnology), HYB331-01 (cat. sc-58749, Santa Cruz Biotechnology), 121-3 (cat. ab270732, Abcam), 35I9 (cat. ab27156, Abcam), rDSD/4565 (cat. ab273137, Abcam), AE-2 (cat. MA1-35346, Thermo Fisher Scientific). Antibodies were tested for binding to dsDNA using a mouse anti-dsDNA IgG ELISA kit per the manufacturer’s instructions (cat. 5120, Alpha Diagnostic International).

### Electrophoretic mobility shift assays (EMSA)

Widom601 dsDNA complexed with recombinant human histones was purchased from Epicypher (cat. 16-0009). Free Widom601 dsDNA was amplified from a purchased template (cat. 18-0005, Epicypher) using primers 5’-CTGGAGAATCCCGGTGC and 5’-ACAGGATGTATATATCTGACACGTGC. Double-stranded DNA (free and/or histone bound) was combined at a final concentration of 4ng/μL total DNA with varying concentrations of 35I9 in PBS (21-040-CM, Corning) in 10μL total. 1 μL of Novex high density TBE sample buffer (cat. LC6678, Thermo Fisher Scientific) was added and 10μL of mixture was loaded into 6% DNA Retardation Gels (cat. EC6365BOX, Thermo Fisher Scientific). Gels were run at 4°C, 100V for 120 minutes in 0.5x TBE, stained with SYBR Safe (cat. S3312, Thermo Fisher Scientific) at 1:10000 dilution in 0.5x TBE for 30 minutes and imaged on an ImageQuant LAS4000.

### Immunoblot analysis

EMSA gels, after imaging for dsDNA with SYBR Safe, were transferred to nitrocellulose membranes using the iBlot2 dry transfer system (Thermo Fisher Scientific) with transfer voltage of 23V for 6 minutes. Membranes were blocked by incubating with 5% milk in tris-buffered saline + 0.1% Tween (TBST-M) for 1 hour at room temperature with gentle rocking. Membranes were then incubated with HRP-conjugated primary antibody against human histone H3 (cat. ab21054, Abcam) at 1:2000 dilution in TBST-M for 2 hours at room temperature. After incubation, membranes were washed with tris-buffered saline + 0.1% Tween (TBST) three times (5 minutes each) with gentle rocking followed by a final 5-minute wash in tris-buffered saline (TBS). Bound antibody was detected with Novex ECL chemiluminescent substrate (cat. WP20005, Thermo Fisher Scientific).

### Transmission Electron Microscopy (TEM) imaging

The morphologies of nucleosomes (NCPs, Epicypher cat. 16-0009), 35I9 anti-DNA mAb (cat. ab27156, Abcam), and NCP-mAb conjugates were confirmed by negative-stained TEM imaging. NCP-mAb were prepared and imaged at different NCP: mAb ratios (1:2 1:1, 2:1,4:1). For sample preparation, first, ultrathin carbon film Au grids (CF300-Au-UL, Electron Microscopy Sciences) were plasma treated using a Denton Sputter Coater at 5 mA for 8 secs. Next, 5 μL of sample solution (20 ng/μL in PBS) was incubated on the plasma-treated grids for 1.5 mins and the excess of liquid was removed with a filter paper (pore size 110 nm, Whatman). The grid was washed twice by dropping 5 μL of deionized water on the grids and quickly removed using a filter paper. Negative staining was performed by dropping 5 μL of 2% uranyl acetate solution on the grid for 18-20 secs and the excess of liquid was removed by a filter paper. Imaging was performed on a JEOL 2100F TEM at 200 kV. All images were recorded on a Gatan UltraScan charge-coupled device (CCD) camera.

### Cell culture

Expi293F cells (cat. A14527, Thermo Fisher Scientific) were maintained in Expi293 Expression Medium (cat. A1435101, Thermo Fisher Scientific) at densities of 0.25-6×10^6^ cells/mL in a humidified atmosphere of 95% air and 8% CO_2_ at 37°C. Luc-MC26 (carrying firefly luciferase, from the Kenneth K. Tanabe Laboratory, Massachusetts General Hospital) were cultured in ATCC-formulated Dulbecco’s Modified Eagle’s Medium DMEM (cat. D6429, Sigma) supplemented with 10% fetal bovine serum (cat. A5256701, Gibco) and 1% penicillin/streptomycin (cat. 30-002-Cl, Corning) in a humidified atmosphere of 95% air and 5% CO_2_ at 37°C.

### Antibody expression and purification

Desired Fc mutations were introduced into the heavy chain sequence (as determined by LC-MS de-novo sequencing, Rapid Novor Inc) and codon-optimized for expression in HEK293 cells. Gene-blocks for the heavy and light chain (IDT) were cloned into the same gWiz plasmid, separated by the T2A ribosome skipping sequence (*65, 66*). Expi293F cells at a density of 3×10^6^ cells/mL were transfected with 1mg/L of culture of plasmid complexed with PEI Max 40K (cat. 24765-100, Polysciences) in a 1:2 plasmid:PEI w/w ratio in 40mL Opti-MEM (cat. 31985062, Thermo Fisher Scientific) per 1L culture. Flasks were kept in a shaking incubator (125rpm) at 37°C and 8% CO_2_. 24h after transfection, flasks were supplemented with glucose and valproic acid (cat. P4543, Millipore Sigma) to final concentrations of 0.4% v/v and 3mM respectively. Culture supernatant was harvested after 5-6 days and purified using Protein A affinity chromatography (AKTA, Cytiva), buffer exchanged into PBS and sterile-filtered.

### Blood collection and processing

Retro-orbital blood draws (70μL in general, 50μL for antibody pharmacokinetic study) were collected via non-heparinized capillary tubes from mice under isofluorane anesthesia, alternating between eyes for serial draws. Mice were allowed to recover from anesthesia between blood draws. Blood was immediately displaced from capillary tube into 70μL of 10mM EDTA (cat. AM9260G, Thermo Fisher Scientific) in PBS. For terminal bleed samples, blood was collected via cardiac puncture into a syringe filled with 200μL of 10mM EDTA in PBS. Total volume was measured and additional 10mM EDTA in PBS was added to reach a 1:1 ratio of blood to EDTA. Blood with EDTA was kept on ice and centrifuged within 90 minutes at 8000g for 5 minutes at 4°C. The plasma fraction was collected and stored at −80°C until further processing.

### DNA extraction

Frozen plasma was thawed and underwent ultracentrifugation at 15,000g for 10 minutes. Sample volumes were normalized to 2.1mL with addition of PBS. DNA was extracted using the QIAsymphony DSP Circulating DNA kit (cat. 937556, Qiagen) on a QIAsymphony instrument (Qiagen) per the manufacturer’s instructions. gDNA was extracted from cell lines or buffy coat using the QIAsymphony DSP DNA Midi Kit (cat. 937855, Qiagen) per the manufacturer’s instructions.

### cfDNA pharmacokinetic assays

10-20ng of W601 sequence (free and histone bound, cat. 16-0009, Epicypher) was combined with antibody in 200μL of PBS. Engineered variants were produced in-house; 35I9 was purchased from Abcam (cat. ab27156); mouse IgG2a control (clone 20102, cat. MAB003), anti-FcγRII/III (rat anti-mouse, clone 190909, cat. MAB1460), and anti-FcγRI (rat anti-mouse, clone 29035, cat. MAB2074) were purchased from R&D systems. 40μg of anti-FcγRII/III and 20μg of anti-FcγRI was used in FcγR-blocking conditions. Mixtures were kept on ice until injection. Each mouse was anesthetized with inhaled isofluorane and injected i.v. with 200μL of mixture. At 1-minute after injection, 70μL of blood was collected via a retro-orbital blood draw. Mice were allowed to recover after this and between subsequent blood-draws (all 70μL). Cell-free DNA was extracted from plasma and W601 levels were quantified using Taqman qPCR (PrimeTime, cat. 1055770, IDT) with the following primers and probes: forward, 5’CGCTCAATTGGTCGTAGACA; reverse, 5’TATCTGACACGTGCCTGGAG; probe, /56-FAM/TCTAGCACC/ZEN/GCTTAAACGCACGTA/3IABkFQ/.

### Antibody pharmacokinetics and biodistribution

To label antibodies, the amine-reactive fluorophore AQuora® 750 (cat. AQ-111960LF, Quanta Biodesign) was incubated with 100μL aliquots of 1.3mg/ml aST3 or 1.0mg/ml 35I9, at 8 dye:1 protein and 18 dye:1 protein molar ratio, respectively, for 2.5h at room temperature. Excess dye was removed by washing conjugate with sterile DPBS 10 times using 30kDa Amicon filters (cat. UFC503024, EMD Millipore) at 12,000rpm, 2.5min, 4°C. After concentrating the volume of the last wash to approximately 200μL, a Nanodrop spectrophotometer (Thermo Fisher Scientific) was used to measure the protein yield (absorbance at 280nm) and the fluorophore concentration and degree of labelling (absorbance at 740nm) to confirm comparable degree of labeling for both proteins (aST3: 2.8 fluorophores/protein and 35I9: 3.1 fluorophores/protein). For antibody pharmacokinetic studies (n=5 per group), AQuora® 750-labeled antibodies were injected i.v. at 4.0mg/kg (200μL in sterile DPBS) into anesthetized mice and 50μL of blood collected from alternating eyes at 1min, 30min, 1h, 2h 6h and 24h. Abundance of antibodies in plasma samples was measured using a Tecan fluorometer 750/785nm and presented as % remaining relative to the 1 min plasma sample. For biodistribution studies (n=5 per group), AQuora® 750-labeled antibodies were injected i.v. at 4.0mg/kg (200μL in sterile DPBS) into anesthetized animals. 1h after injection, mice were euthanized via isoflurane overdose and perfused with 30mL PBS to remove blood from circulation. Liver, kidneys, spleen, heart, and lungs were harvested and imaged using an Odyssey DLx imaging system (Licor). AQuora® 750-fluorescence values were normalized to organ surface area in the imaging scan as a surrogate of organ weight.

### Animal models

All animal studies were approved by the Massachusetts Institute of Technology Committee on Animal Care (MIT Protocol 0420-023-23). Animals were maintained in the Koch Cancer Institute animal facility with a 12h-light/12h-dark cycle at 18-23°C and 50% humidity. All animals received humane care, and all experiments were conducted in compliance with institutional and national guidelines and supervised by staff from the Division of Comparative Medicine of the Massachusetts Institute of Technology. To generate the lung metastasis model, 1×10^6^ Luc-MC26 cells in 100μL DPBS were injected i.v. into female Balb/c mice (4-6 weeks, Taconic Biosciences) and allowed to grow for 2 weeks. Tumor growth was monitored longitudinally via intraperitoneal injection of luciferin and luminescence imaging using the In Vivo Imaging System (IVIS, PerkinElmer). Tumor burden at end point was measured via ex vivo luminescence imaging of lungs in the IVIS system.

### Assessing the performance of DNA-binding agent for tumor detection

Between days 10 and 12 post-tumor inoculation, the performance of aST3 on ctDNA testing was assessed in Luc-MC26-tumor bearing mice. As an internal control, 70μL blood was sampled retro-orbitally from each mouse prior to treatment. Subsequently, 4.0mg/kg aST3 (in 200μL sterile DPBS) or 4.0mg/kg IgG2a isotype were administered into awake mice i.v. 2h after treatment, 70μL of blood was collected retro-orbitally from the contralateral eye and mice exsanguinated via cardiac puncture to collect the remainder of blood for analysis. cfDNA concentration measurement and ctDNA detection was performed on all samples as described below.

### Cell-line and buffy coat sequencing and fingerprint design

gDNA was extracted from Luc-MC26 and Balb/c buffy coat, sheared, and libraries were prepared using the Kapa HyperPrep Library Construction kit (cat. KK8504, Roche Diagnostics). Whole genome sequencing was performed to 30x coverage for Luc-MC26 and 15x coverage for Balb/c buffy coat. Sequencing data was aligned against mm9 genome assembly, mutation calling with Mutect2, and tumor SNV fingerprint selection for a 2000-SNV panel as previously described (*6*). A pool of 120-bp biotinylated probes against the 2000 sites was ordered from IDT. 1,822 sites validated against Luc-MC26 gDNA.

### cfDNA sequencing and mutation detection

Library construction for cfDNA was conducted using custom adapters with UMIs to enable duplex sequencing. Two rounds of hybrid capture were performed followed by sequencing, alignment to mm9, duplex consensus calling, and identification of mutation-bearing duplexes (“mutant molecules”) using our group’s established workflow as previously described (*6*).

### Sensitivity estimation

To estimate sensitivity at smaller panel sizes, we employed a bootstrap procedure downsampling with replacement from our 2000-site panel to smaller panel sizes. Sensitivity at different detection thresholds was estimated as the fraction of mice that had mutant molecules detected at the given threshold. For each panel size and dose, 100 replicates were generated, and the mean sensitivity and standard error was computed. To estimate sensitivity at lower tumor fractions, we first confirmed that the distribution of mutant molecules (*n_ij_* and the distribution of the ratio of mutant molecules to total molecules (*n_ij_/t_ij_*) could be accurately recapitulated via a binomial sample *n_ij_* ~ Binom(*t_ij_, f_i_*) where *n_ij_* is the number of mutant molecules at site *j* in sample *i, t_ij_* is the number of total molecules at site *j* in sample *i*, and *f*_i_ is the global tumor fraction in sample *i*. To estimate sensitivity at lower tumor fractions, we then generated distributions of mutant molecules under lower *f_i_* for each sample, also incorporating various panel sizes as above, and computed sensitivity for detection of mutant molecules under various detection thresholds. Sensitivity at each *f_i_*, dose, and panel size was estimated by taking the mean and standard error from 100 replicates.

### Identification of enriched sites

The duplex depth within each dose level was normalized to mean 0 and standard deviation 1. For each site, the Pearson correlation between aST3 dose and normalized duplex depth was computed (with the control antibody group taken as dose = 0mg/kg), and sites were ordered based on the correlation measurement to identify those with relative enrichment in presence of higher antibody doses.

### Analysis of overlap with DNase HS and CpG sites

We downloaded DNase HS peaks for all leukocyte and myeloid datasets in mouse ENCODE (ENCFF063EHX, ENCFF125IXR, ENCFF171XTE, ENCFF185ZCW, ENCFF215CPX, ENCFF359GEV, ENCFF434GOV, ENCFF550NKM, ENCFF566TDU, ENCFF689PKR, ENCFF702OKE, ENCFF754XGR, ENCFF761OVL) and converted to mm9 coordinates using LiftOver (https://genome.ucsc.edu/cgi-bin/hgLiftOver). ENCSR000CMQ and ENCSR000CNP were excluded due to very low read depth. Distance between each of the 2000 sites and the nearest DNase HS peak was computed, with a distance of 0 indicating overlap between the site and a peak. Coordinates of mouse CpG islands were downloaded from UCSC genome browser (table cpgIslandExt, assembly mm9). GC content of each site was computed as the percentage of G/C bases in the 120bp probe sequence.

### DNase protection assays

To measure sensitivity to DNase digestion, the DNaseAlert kit (cat. 11-02-01-04, IDT) was used in combination with various concentrations of recombinant DNase I and antibody 35I9 in 100μL reactions incubated at 37°C in a Tecan microplate-reader with initial measurement before addition of DNase I and subsequent measurements every 5 minutes after addition of DNase I (excitation 365nm, emission 556nm).

### Statistical analysis

One-way ANOVA was used for statistical testing unless noted otherwise. A suite of scripts (Miredas) was used for calling mutations and creating metrics files (*6, 67*). All analysis was performed using custom Python scripts and R (v4.0.3).

## Acknowledgments

We thank the Koch Institute’s Robert A. Swanson (1969) Biotechnology Center for the technical support, and specifically the Nanotechnology Materials Core (RRID:SCR_018674) for the support on TEM imaging. We thank Sarah Cowles, Emi Lutz, and K. Dane Wittrup for advice and assistance with mammalian expression. We thank Greg Gydush for his help setting up the data analysis pipeline used in this work and Leslie Gaffney for her assistance in designing the graphics for this work.

## Funding

Koch Institute Frontier Research Program via the Casey and Family Foundation Cancer Research Fund (SNB, JCL)

Bridge Project, a partnership between the Koch Institute for Integrative Cancer Research at MIT and the Dana-Farber/Harvard Cancer Center (VAA, JCL)

ASCO Conquer Cancer Foundation Young Investigator Award (2021YIA-5688173400) (ST)

Prostate Cancer Foundation Young Investigator Award (21YOUN20) (ST)

“La Caixa” Foundation (ID 100010434), fellowship code LCF/BQ/AA19/11720039 (CMA)

Ludwig Center at MIT’s Koch Institute (CMA)

Koch Institute’s Marble Center for Cancer Nanomedicine (CMA)

T32 (T32HL116275) (SP)

Cancer Center Support (Core) Grant P30-CA14051 from the National Cancer Institute

Gerstner Family Foundation

SNB is a Howard Hughes Medical Institute Investigator.

## Author contributions

Conceptualization: ST, CMA, JCL, VAA, SNB

Methodology: ST, CMA, KX, TB, SS, ZA, SP, CAN, SRA, STW, DS

Investigation: ST, CMA, KX, TB, SS, ZA, SP, CAN, SRA, STW, DS

Visualization: ST, CMA

Funding acquisition: SNB, VAA, JCL, TRG

Project administration: ST, CMA

Supervision: SNB, VAA, JCL

Writing – original draft: ST, CMA

Writing – review & editing: ST, CMA, SNB, VAA, JCL

## Data and materials availability

All sequencing data generated in this study will be deposited into a controlled access database.

## Supplementary Materials

Figs. S1 to S10

Data S1

