## Supplementary Figures for "An intravenous DNA-binding priming agent protects cell-free DNA and improves the sensitivity of liquid biopsies"

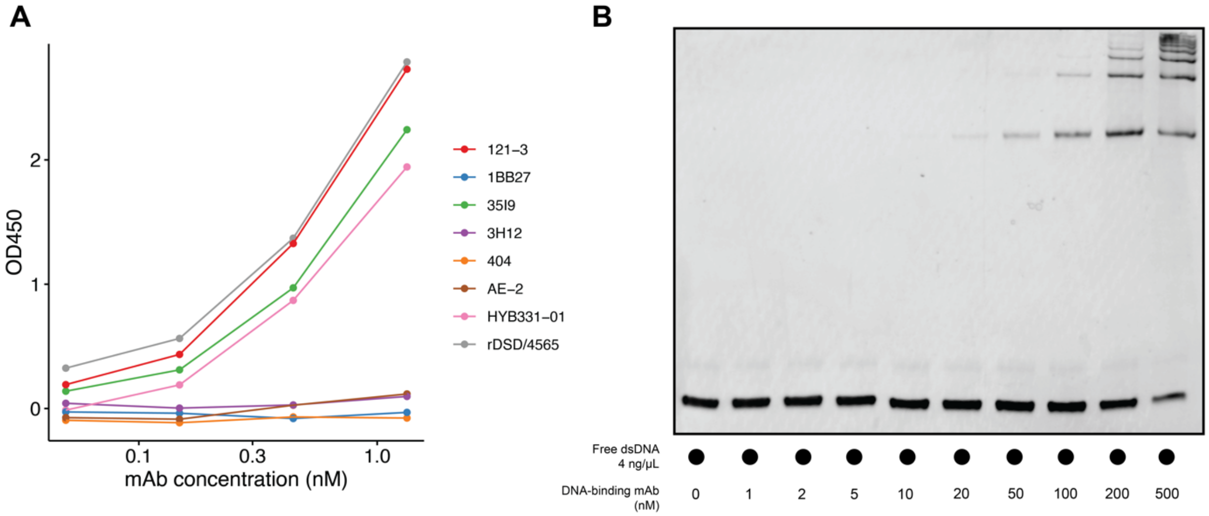


**Fig. S1. Binding of anti-DNA mouse IgG mAbs to free dsDNA. A**, dsDNA binding activity via ELISA for nine DNA-binding mouse IgG antibodies. **B**, EMSA of free W601 dsDNA with DNA-binding mAb 35I9.

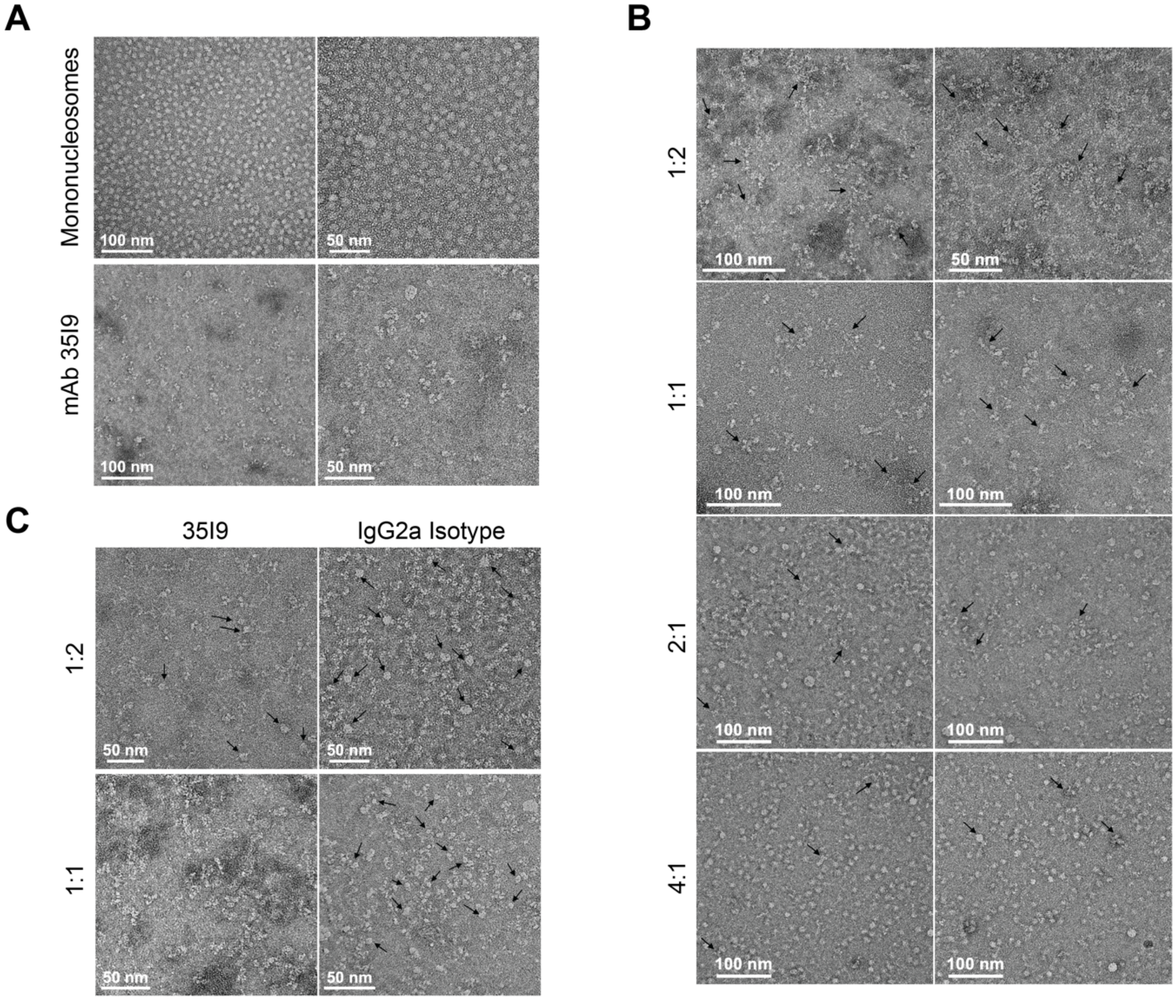


**Fig. S2. TEM images of interaction of mAb 35I9 with mononucleosomes. (A)** Negative stained TEM images of mononucleosomes (NCPs) and anti-DNA mAb 35I9 presented at two magnifications. NCPs have average diameters of 11.4 ± 1.3 nm (81 particles) measured by the ImageJ software. Total protein concentration in each sample is 17 ng/µL. **(B)** Negative stained TEM images of NCP- 35I9 mAb conjugates at different NCP: mAb ratios. Two images from the same samples are presented. Increased aggregation and reduced contrast of intact NCPs were observed at higher mAb concentrations, which agreed with the NCP-mAb interactions observed in electrophoretic mobility shift assays. The black arrows are pointed to additional contrasts from NCP or free DNA at the mAbs. Total protein concentration in each sample is 20 ng/µL. **(C)** Qualitative comparison of the negative stained TEM images of NCPs in the presence of mAb and IgG2a isotype at NCP: Ab ratios of 1:1 and 1:2. More intact NCPs (black arrows) were observed in the isotype samples, confirming a higher affinity of anti-DNA mAb to NCPs. Total protein concentration in each sample is 20 ng/µL.

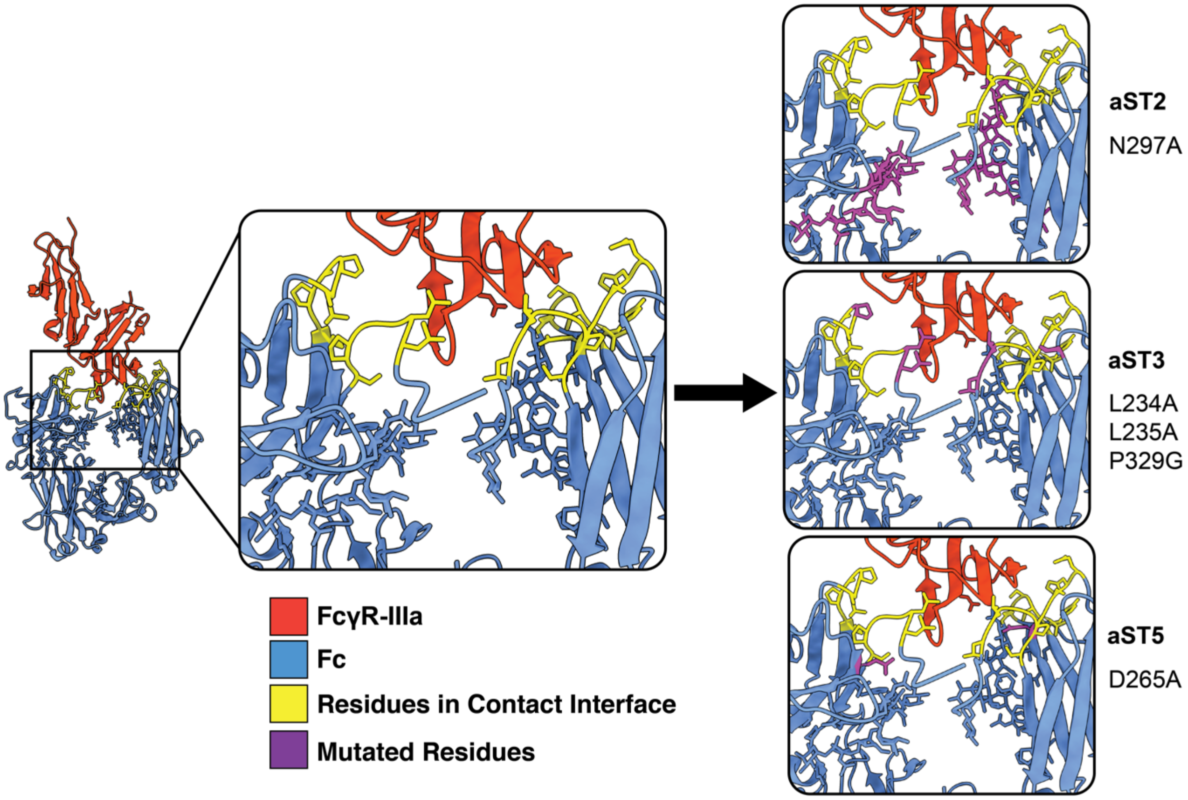


**Fig. S3. Mutations disrupting interaction of Fc with FcγR-IIIa.** Crystal structure of human FcγR-IIIa complexed with human IgG1 Fc (PDB: 1E4K), with residues in contact interface as identified in (*68*) highlighted in yellow. Residues mutated in Fc-engineered variants are conserved between mouse IgG2a and human IgG1, all overlap with the contact interface, and are highlighted here in purple.

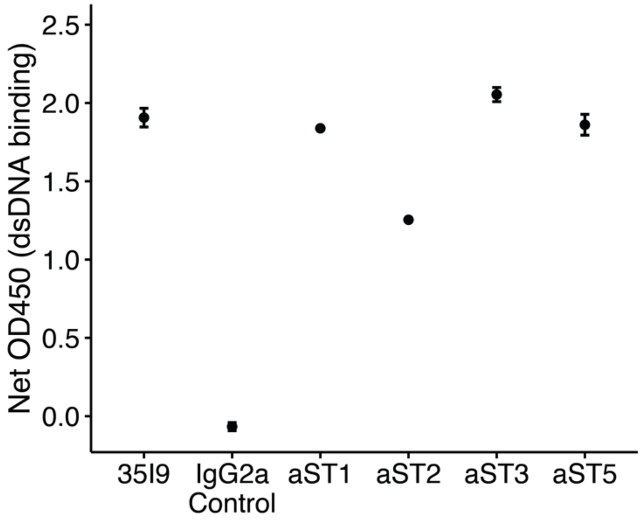


**Fig. S4. Binding of engineered anti-DNA mAbs to dsDNA.** dsDNA binding activity via ELISA for 35I9, unrelated IgG2a mAb, and four engineered variants. All antibodies are at 1nM.

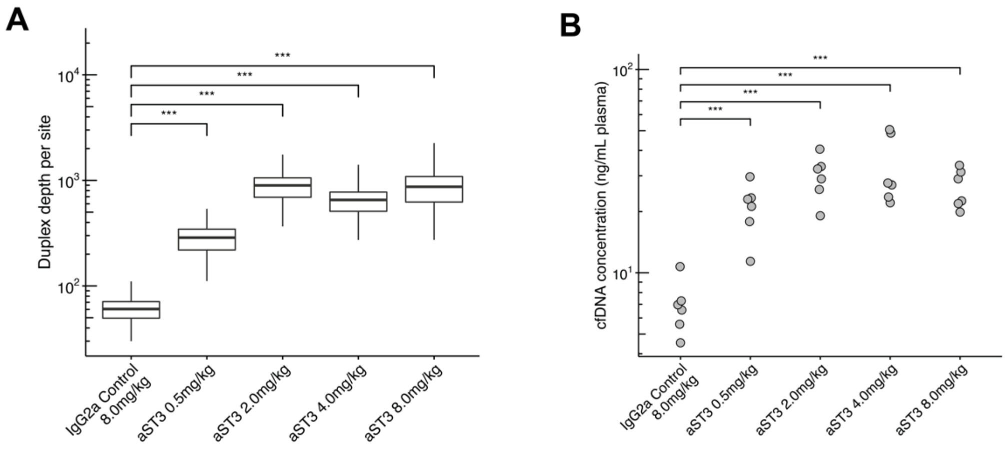


**Fig. S5. Duplex depth and cfDNA concentration improve with priming. (A)** Per-site duplex depth, n=1822 sites x 6 mice per group. (**B**) cfDNA concentration, n=6 mice per group. **A, B** *** P < 0.001; one-way ANOVA.

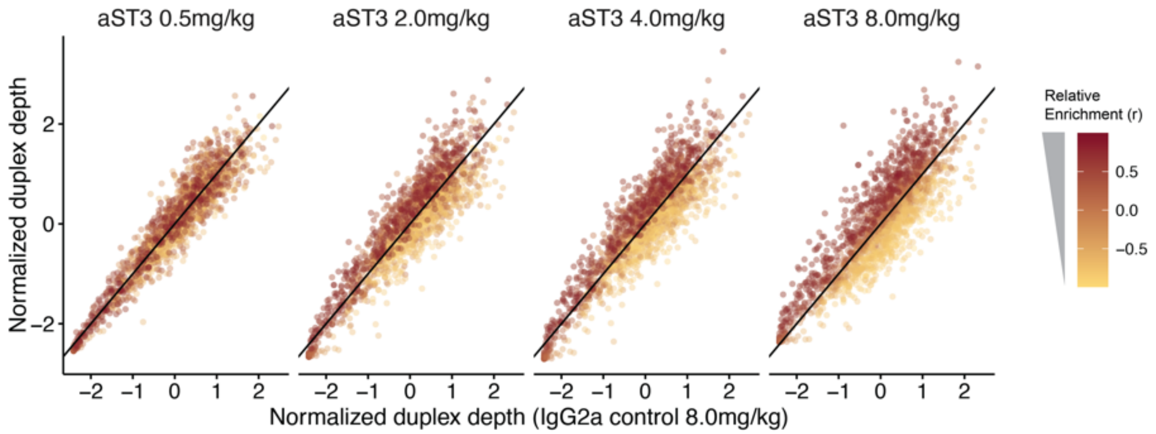


**Fig. S6. Normalized duplex depth in each dose group compared to the control group.** Normalized duplex depth at each of 1,822 sites in each aST3 dose group compared to the IgG2a control group. Each site is colored by correlation between aST3 dose and duplex depth for that site. Black line is the identity (y = x) line.

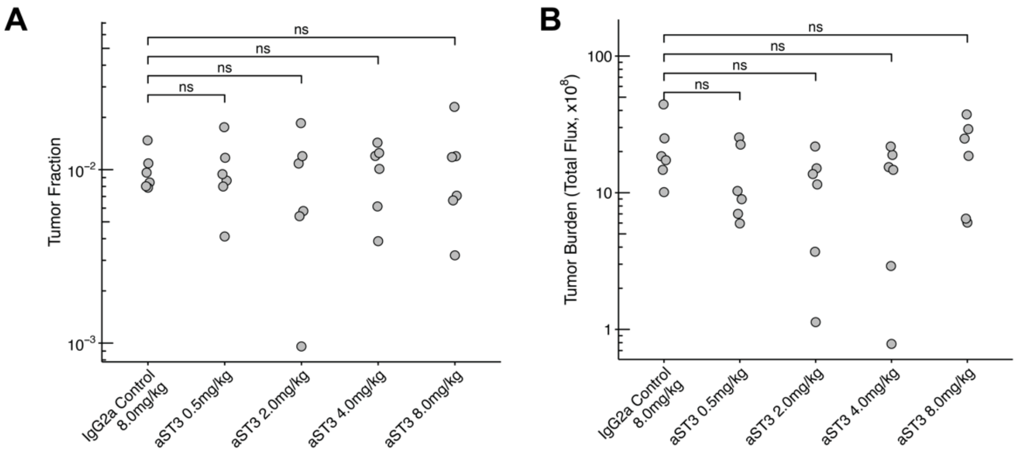


**Fig. S7. Tumor fraction and tumor burden are comparable between groups. (A)** Pre-injection tumor fractions for the five treatment groups. **(B)** Ex vivo lung tumor burden as measured by bioluminescence. **A,B** n=6 mice per group, ns=not significant (P > 0.05), one-way ANOVA.

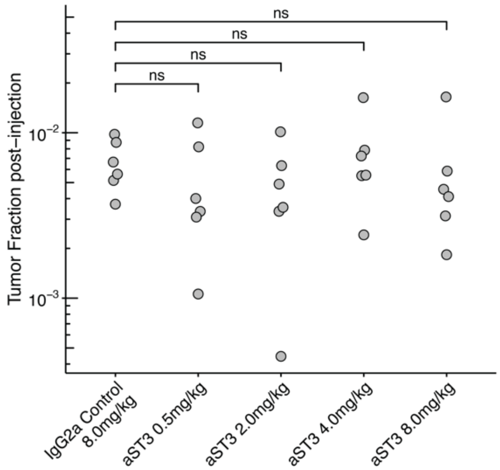


**Fig. S8. Post-injection tumor fraction is comparable between groups.** Post-injection tumor fractions for the five treatment groups, n=6 mice per group. ns=not significant (P > 0.05), one-way ANOVA.

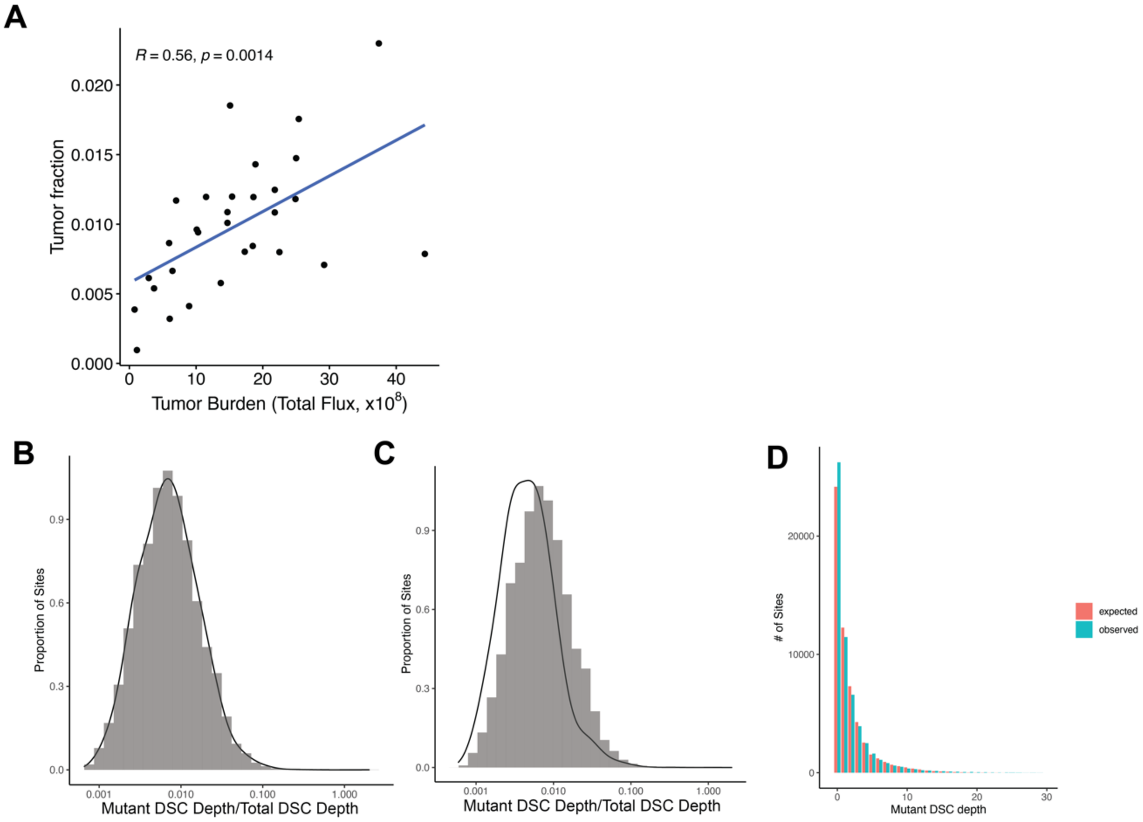


**Fig. S9. A binomial model recapitulates distribution of mutant molecules. (A)** Correlation between lung tumor burden and concentration of mutant molecules per unit volume of plasma. **(B)** Comparison of resampled mutant duplex sequence consensus (DSC) depth/total DSC depth using a binomial model and the per-sample global tumor fraction (black density line) versus the observed distribution (bars). **(C)** Same as **(B)**, but with per-sample global tumor fraction halved. **(D)** Observed distribution of mutant DSC depth versus expected distribution using a binomial model.

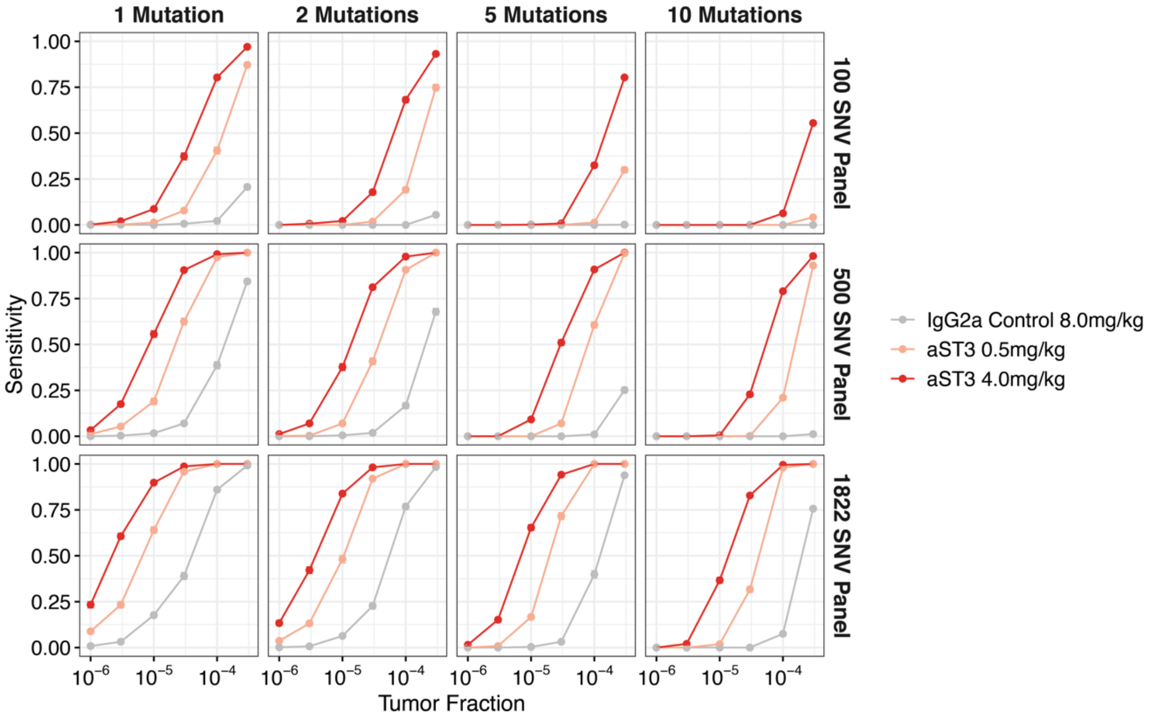


**Fig. S10: Sensitivity for ctDNA detection with various panel sizes and detection thresholds.** Estimated sensitivity at various detection thresholds (1-10 mutations) and panel sizes (100-1822 SNV panels) with and without priming. Points and error-bars represent mean and s.e.m. from 100 replicates.
